# Lung Immune Tone Regulation by the Gut-Lung Immune Axis: Short-chain Fatty Acid Receptors FFAR2 and FFAR3, and IL-1β Expression Profiling in Mouse and Human Lung

**DOI:** 10.1101/2020.08.28.213439

**Authors:** Qing Liu, Xiaoli Tian, Daisuke Maruyama, Mehrdad Arjomandi, Arun Prakash

## Abstract

Microbial metabolites produced by the gut microbiome, such as short-chain fatty acids (SCFA), can influence both local intestinal and distant lung physiology and response to injury. However, how lung immune activity is regulated by SCFAs is unknown. We examined fresh human lung tissue and observed the presence of SCFAs with large inter-individual and even intra-lobe variability. *In vitro*, SCFAs were capable of modifying the metabolic programming in both resting and LPS-exposed alveolar macrophages (AM). Additionally, since we hypothesized that lung immune tone could be defined through priming of the inflammasome (aka signal 1), we interrogated naïve mouse lungs for pro-IL-1β message and localized its presence within the alveolar space *in situ*, specifically in AM subsets, and in close proximity to alveolar type 2 epithelial (AT2) cells. We established that metabolically active gut microbiota, that produce SCFAs, can transmit LPS and SCFAs to the lung (potential sources of signal 1), and thereby could regulate lung immune tone and metabolic programming. To understand how murine lung cells sensed and upregulated IL-1β in response to gut-microbiome factors, we determined that *in vitro*, AM and AT2 cells expressed SCFA receptors, FFAR2, FFAR3, and IL-1β but with different expression patterns and LPS-inducibility. Finally, we observed that IL-1β, FFAR2 and FFAR3 were expressed both in isolated human AM and AT2 cells *ex-vivo*, but in fresh human lung sections *in situ*, only AM expressed IL-1β at rest and after LPS challenge. Together, this translational study using mouse and human lung tissue and cells supports an important role for the gut microbiome and SCFAs in regulating lung immune tone.

## INTRODUCTION

The human gut and lung microbiomes have been linked to host health and immune development (14, 22, 29, 32, 45). Key microbial metabolites such as short-chain fatty acids (SCFA), which originate via microbiome fermentation or from the diet, have been shown to regulate local tissue inflammation, and act as communication signals to extra-intestinal organs (28, 43, 46). SCFAs can influence allergic and inflammatory lung diseases including asthma (16, 21, 50, 53). However, the lung microbiome’s low biomass (10^3^-10^5^/g) (55) and lack of significant metabolic activity imply that they are an unlikely source of SCFA production. Consequently, we and other investigators have proposed that SCFAs modulate lung immune function via direct or indirect mechanisms (7, 11, 50), but thus far no consensus has been established on how gut-originating SCFAs regulate lung inflammatory responses.

The lung likely exists in a state of immune responsiveness that is influenced by numerous factors, including environmental exposures, genetics, diet, medications, prior disease/injury exposure, and microbiome. Many of these factors are complexly intertwined and likely contribute to a dynamic *‘lung immune tone’* that informs the host lung response to the next injury or infectious exposure. Microbiome-sourced bacterial signatures, such as LPS, and metabolites, such as SCFAs, could contribute to creating this lung immune tone within the gut-lung immune axis. The primary metabolite receptors for SCFAs are two free fatty acid receptors (FFAR): FFAR2 (GPR43) and FFAR3 (GPR 41) (6, 34, 37, 40). FFAR2 is engaged primarily by acetate and propionate, and FFAR3 by propionate and butyrate (25, 47). Expression of these FFARs has been described in the intestinal stromal and immune cells, mainly in the colon, but FFAR2 and FFAR3 have also been detected extra-intestinally (5, 28, 34).

We previously demonstrated that sensing of LPS and SCFAs produced by the gut microbiome could strongly influence the course of lung injury and infections, thereby supporting the existence of the gut-lung axis of immune communication (41, 50). Moreover, we reported that co-exposure of bacterial products, such as LPS, with SCFAs enhanced or repressed inflammatory responses depending on their concentrations or levels within the lung (50) (reflecting the presence of specific types of gram-negative and gram-positive fermenting bacteria in the microbiome). We further reported low but detectable resting IL-1β expression in lung tissue and interpreted this as gut-microbiome mediated inflammasome priming (signal 1) (50). The inflammasome function was also required for appropriate host response to lung injury and infection (51).

In the present study we characterized the pulmonary component of the gut-lung immune axis and elucidated how SCFA and microbial product presence and sensing in the lung may contribute to steady-state lung immune tone. To do so, we assessed the expression and regulation of FFAR2 and FFAR3 using mouse cell lines and freshly isolated and cultured human lung cells. We also examined intact mouse or human lung tissue to identify the cellular source(s) of IL-1β, FFAR2 and FFAR3 expression. By identifying lung cell types that receive the gut-derived signals (LPS, SCFAs, etc.) and thereby establish lung baseline IL-1β expression, we hope to advance our understanding of the potential interactions between the gastrointestinal and pulmonary systems in health and disease.

## MATERIALS AND METHODS

### Animal care

All studies were approved by the institutional animal care and use committee at University of California, San Francisco (Protocol# AN152943). All mice were purchased (The Jackson Laboratory, Bar Harbor, ME) or bred in the animal facility at University of California, San Francisco. Wild-type (WT) C57BL/6, C3H/HeOuJ mice, and germ-free (GF aka gnotobiotic) mice were used in this study (8-10 weeks old GF or 10-15 weeks old WT C57BL/6J and C3H/HeOuJ adult male mice). GF mice were obtained from the UCSF Gnotobiotic Facility. In accordance with Biosafety Level 2 guidelines from the Centers for Disease Control and Prevention, GF mice were maintained in sterile gnotobiotic isolators under a strict 12hr light cycle and were fed an autoclaved standard diet. Commercially purchased mice were allowed to acclimatize to their new housing for at least 1 week before any experiments were conducted on them.

### Human lung tissue samples

Lung tissue samples were collected through an associated lung cancer screening and treatment program, where non-cancerous (“healthy”) portions of the resected lung tissue were excised for this purpose from patients undergoing surgery as part of their definitive lung cancer treatment. The UCSF Institutional Review Board approved the protocols and performance of this study, and written informed consent and Health Insurance Portability and Accountability Act (HIPAA) was obtained from all patients. For metabolomic analyses, lung tissue was immediately frozen on dry ice and stored for later submission for metabolomic analyses (see below). Lung tissue for RNAScope was either fixed right away in formalin or equivalent sections were incubated in PBS or LPS for 2h prior to fixation (see RNAScope section below for more details for subsequent steps).

Human rejected donor lungs were obtained with explicit approval for the use of donor lungs for research from each donor’s family by Donor Network West. All donor lung tissue originated from the right upper lobe of the rejected lungs.

### Reagents and cell lines

Short-chain fatty acids (SCFAs) (acetate, butyrate, propionate), Lipopolysaccharide (LPS), were purchased from Sigma-Aldrich, St. Louis, MO, United States. Seahorse XF calibrant solution (part number 100840-000), Seahorse XF RPMI medium, pH 7.4, (part number 103576-100), and Seahorse XF cell mito stress test kit (part number 103015-100, contains 1 each of oligomycin, FCCP, and rotenone/antimycin A. Oligomycin, FCCP, antimycin A and rotenone) all purchased from Agilent Technologies, Inc.

The following cell lines were used in this study: MLE 12 (mouse lung epithelial AT2), MH-S (wild-type BALB/c alveolar macrophage cell line, ATCC), EOMA (129 background endothelial cell line, ATCC, Manassas, VA, United States), HUVEC (primary human umbilical vein endothelial cells, used at passage 6 or less, Promocell, Heidelberg, Germany), HPMEC-M or HPMEC-F (primary human pulmonary microvascular endothelial cells-male or female, used at passage 2–6, Promocell, Heidelberg, Germany). Human lung alveolar macrophages (AM) were collected by bronchoalveolar lavage (BAL) and alveolar type II (AT2) cells were isolated from elastase-digested human lung tissue using negative selection (anti-CD14 and IgG antibodies) (15). All cells were incubated at 37°C under humidified 5% CO2 and then treated with 10ng/mL or 100ng/mL LPS for 24h or fresh cells were used.

### Bacteroides thetaiotaomicron (*B. theta*) strains inoculation experiment

Two *B. theta* strains: *B.theta* Delta TDK (WT) and *B.theta* Delta 1686-89 (del propionate) strains were a kind gift from Eric Martens, University of Michigan (24, 30). Both *B. theta* strains were grown in brain-heart infusion (BHI) broth in an anerobic chamber. Both *B. theta* strains were used to gavage the GF adult (gnotobiotic) C57BL/6 mice and each individual mouse was colonized using 100ul of a bacterial suspension with each mouse receiving effectively ~10^9^ CFUs. Two weeks later of inoculation, samples of lung tissue, fecal pellets and cecal contents were collected and were homogenized and serially diluted used for endotoxin content measurement.

### Endotoxin measurement

Limulus Amebocyte Lysate (LAL) test cartridges (Endosafe® nexgen-PTS™ Charles River Laboratories) – a rapid, point-of-use handheld spectrophotometer that is used for real-time endotoxin testing – was used in this study for mice lung tissue and stool/intestinal tissue sample endotoxin measurement.

### SCFA and Amino Acid Measurement

Short chain fatty acids of human lung samples were quantified using a Water Acquity uPLC System with a Photodiode Array Detector and an autosampler (192 sample capacity) at the Baylor University Metabolomic Core. Samples were analyzed on HSS T3 1.8 mm 2.1 150 mm column. Flow rate: 0.25 mL/min, injection volume: 5 mL, run-time: 25 min per sample. Eluent A was 100 mM sodium phosphate monobasic, pH 2.5, eluent B was methanol, the weak needle wash was 0.1% formic acid in water, the strong needle wash was 0.1% formic acid in acetonitrile, and the seal wash was 10% acetonitrile in water. The gradient was 100% eluent A for 5 min, gradient to 70% eluent B from 5 to 22 min, and then 100% eluent A for 3 min. The photodiode array was set to read absorbance at 215 nm with 4.8 nm resolution. Samples were quantified against standard curves of at least five points run in triplicate. Standard curves were run at the beginning and end of each metabolomics run. Quality control checks (blanks and standards) were run every eight samples. Results were rejected if the standards deviate by greater than 5%. Concentrations in the samples were calculated as the measured concentration minus the concentration of the solvent; the range of detection was at least 1 – 100 mmol/g stool. Stool samples were similarly analyzed for SCFA levels. Some samples were also sent independently to University of Michigan Metabolomic core to confirm findings and representative data is shown.

### Seahorse™ Extracellular Flux Metabolism Analysis

To examine metabolic response in the MH-S alveolar macrophage (AM) cell lines exposed to LPS and SCFAs, Agilent Seahorse XFe24 analyzer platform (Agilent Technologies, Inc.) was used per manufacturer’s instructions. Briefly, MH-S cells were seeded into XFe24 cell culture plate, 40K/well were grown in RPMI medium supplemented with 10% FBS and 1% P/S. All treatments given 4h after seeding cells. As noted, cells were treated with combinations of LPS (10ng/mL), acetate (5mM), propionate (0.1mM or 5mM).

### RNA Scope™ *In Situ* Hybridization for RNA transcript detection

Lungs from naïve wild-type C3H mice or mice that underwent LPS challenge (10mg/kg IV for 1 h) (41, 51) were fixed in formalin. Human lung tissue was obtained following a lobectomy for tumor resection; the section studied was the furthest section away from the isolated nodule and thus reflective of non-tumor/”normal” lung. Lung sections were either fixed in formalin immediately (fresh) or incubated in PBS or 10μg/mL LPS solution for 2h before fixing. Paraffin embedded sections were stained using RNAScope® (from Advanced Cell Diagnostics – ACDBio) using specific human or mouse RNA probes for IL-1β, IL-18, and for cell markers, such as hopx (AT1 cells), surfactant C (AT2 cells), CD11c (AM and dendritic cells), CD45 (hematopoietic cells), CD206/MRC1 (macrophages). Probes were visualized as two colorimetric dyes (dots or clusters) via light microscopy (mouse) or three fluorescent dyes via fluorescent microscopy (human). Briefly, 5 μm formalin fixed, paraffin-embedded tissue sections were pretreated with heat and protease prior to hybridization with the target oligo probes. Preamplifier, amplifier and AP-labeled oligos were then hybridized sequentially, followed by chromogenic precipitate development. Each sample was quality controlled for RNA integrity with a positive control probe specific to the housekeeping gene peptidylprolyl isomerase B and for background with a negative control probe specific to bacterial Bacillus subtilis gene dihydrodipicolinate reductase (dapB). Optimization of pretreatment conditions was performed to establish the maximum signal to noise ration. Specific RNA staining signal was identified as black, red, or blue-green punctate dots. Samples were counterstained with Gill’s Hematoxylin with nuclei appearing light purple or DAPI.

### Quantitative reverse transcription real-time polymerase chain reaction

TaqMan-specific inventoried gene primers for glyceraldehyde 3-phosphate dehydrogenase (GAPDH), beta actin, β-glucuronidase (GUSB), interleukin (IL)-6, IL-1β, chemokine (C-X-C motif) ligand (CXCL) 1, CXCL2, free fatty acid receptor (FFAR) 2, FFAR3, and toll like receptor (TLR) 4 were used to measure the message levels of these human or mouse genes in cells or lung tissue (Life Technologies, Carlsbad, CA).

Lung tissue was homogenized (Tissue-Tearor - Biospec Products, Bartlesville, OK) and total RNA isolated using Trizol^®^. We used the High Capacity RNA-to-cDNA reverse transcription Kit using 1 μg messenger RNA per reaction (Life Technologies, Carlsbad, CA). Quantitative real-time polymerase chain reaction was performed using the ABI Prism 7000 Sequence Detection System (Life Technologies, Carlsbad, CA). Run method: Polymerase chain reaction activation at 95°C for 20s was followed by 40 cycles of 1s at 95°C and 20s at 60°C.

The average threshold counts (Ct) value of 2–3 technical replicates were used in all calculations. The average Ct values of the internal controls (glyceraldehyde 3-phosphate dehydrogenase, beta actin) was used to calculate ΔCt values for the array samples. Data analysis was performed using the 2^−ΔΔCt^ method, and the data were corrected for statistical analysis using log transformation, mean centering, and autoscaling.

### Sandwich enzyme-linked immunosorbant assay (ELISA)

Concentrations of IL-6 in cell culture supernatant were determined using the mouse DuoSet kit (R&D Systems, Minneapolis, MN). All assays were performed according the manufacturer’s supplied protocol. Standard curves were generated and used to determine the concentrations of individual cytokines or chemokines in the sample. Results are presented as Mean ± SD.

### Statistical Analysis

Data in the figures are expressed as mean +/− SD. Data from *in vivo* studies comparing two conditions were analyzed using 2-tailed nonparametric Mann–Whitney analyses. Data from *in vitro* studies comparing two conditions (*ex vivo* pooled alveolar macrophage stimulation studies) were analyzed using standard Student’s t-test with equal SD to generate *P* values. For analyses of multiple groups (such as for the time course experiments in Figure 2), simple 1-way ANOVA was used and Tukey’s correction for multiple comparisons applied. GraphPad Prism was used for statistical analyses (GraphPad Software, La Jolla, CA). For all *in vivo* experiments, exact *P* values are reported, and for *in vitro* analyses, p values < 0.05 were considered significant. P values are represented as follows in the figures: *< 0.05; **< 0.01; ***< 0.001; ****< 0.0001. Experiments were repeated 2 or more times, as indicated in the figure legends. If any samples were excluded from analysis, the number of samples excluded and the reasons for exclusion are included in the figure legends for the corresponding figures.

## RESULTS

### Human lung tissue contains variable micromolar levels of acetate and propionate

Human lung tissue from 5 patients undergoing lung lobectomies were assayed SCFA (including medium and branch-chain fatty acids: C2-C8) levels. We noted significant variability in the relative levels of all SCFAs between patients (**Figure 1A**) as well as when comparing the absolute levels of the SCFAs (**Figure 1B**, left) or as a percentage of total SCFAs (**Figure 1C**, left). Previously, we had reported that the ratios of C2:C3 can regulate lung inflammatory responses (50) and these varied from 3:1 to 10:1 in this cohort (**Figure 1B**, right and **Figure 1C**, right).

**Figure 1.**
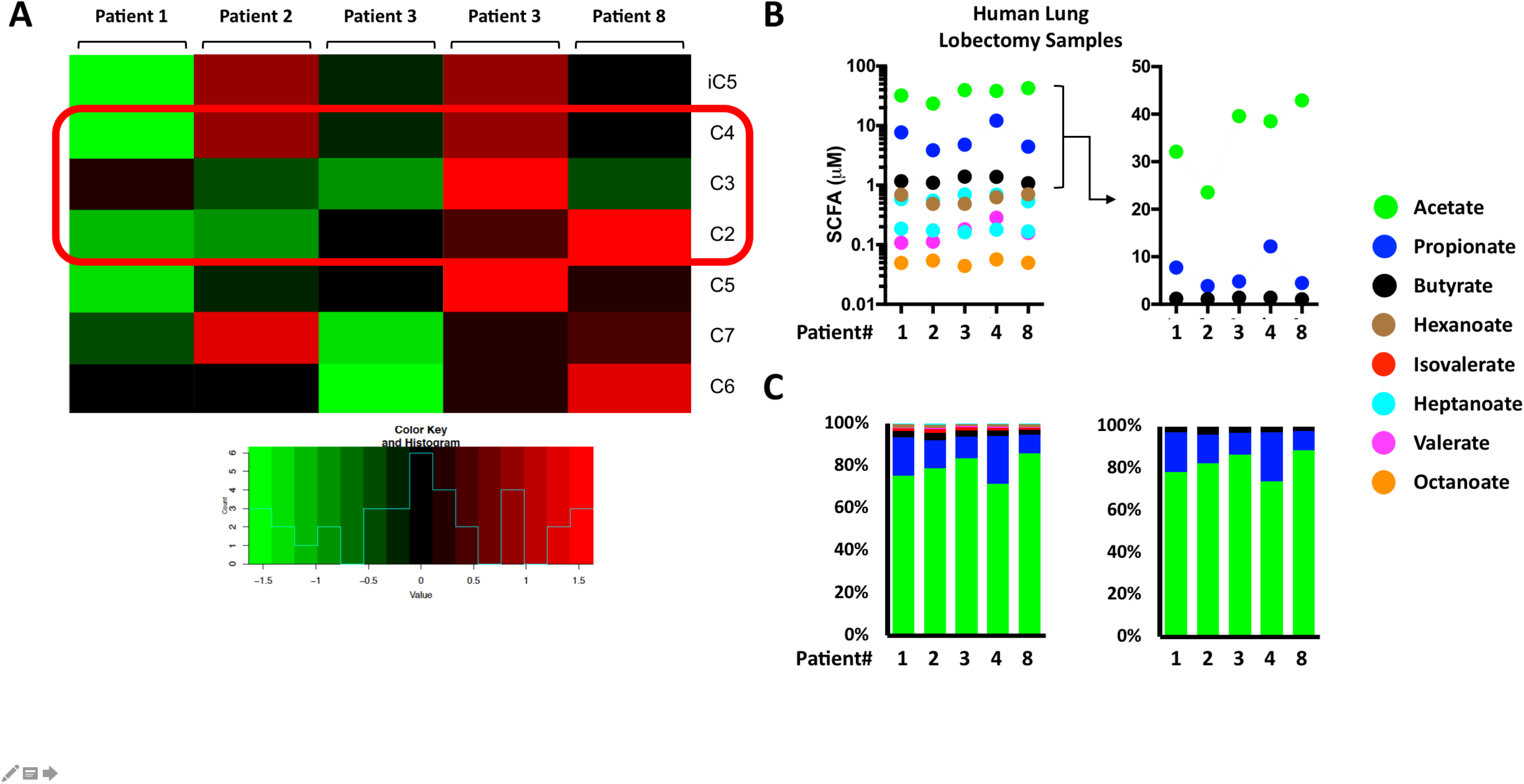
Large variability exists in SCFA levels in human lung tissue from different patients and even within a human lung single lobe. (A) Relative levels of SCFAs from human lung tissue (n = 5 patients: Patient 0001, 0002, 0003, 0004, and 0008) obtained from surgical resections (lung lobectomies) for isolated tumor nodules. (B) Absolute concentrations of SCFAs in each of the lung sections (left) and focusing only on C2-4 (right). (C) Presented as a percentage of total SCFAs (all SCFAs, left and only C2-4, right). SCFAs (C2-4) are highlighted by the red box. SCFA = short chain fatty acid, C2 = acetate, C3 = propionate, C4 = butyrate, iC4 = isobutyrate, C5 = valerate, iC5 = isovalerate, C6 = hexanoate, iC6 = isopropyl hexanoate, C7 = heptanoate, C8 = octanoate, RUL = Right Upper Lobe.

### LPS and SCFA presence in naïve murine lungs depends on the presence of an intact gut microbiome

To confirm that the gut microbiome was the source of lung SCFAs, we examined the metabolomic profile of colons/stool and lung tissue of mice that originated from UCSF’s germ-free (GF aka gnotobiotic) and SPF (specific pathogen free, conventionally housed) facilities. We observed that GF mice had low to absent levels of SCFAs (C2-8) confirming wide-spread literature reports that SCFAs are not generated in mammalian cells and are solely derived from metabolically-active and/or fiber-fermenting gut bacterial species (**Figure 2A**). We measured the levels of LPS present in GF versus SPF mouse stool and lung tissue and noted the presence of LPS in the lungs of SPF mice compared to GF mice (**Figure 2B**, **right**). As a proof of principle, we mono-colonized GF mice with a gram-negative bacterial species (*B.theta*) and found that this colonization led to an increase in LPS within the intestine compared to GF mice; as well as a large increase in LPS levels (albeit extremely low absolute LPS concentrations) in the lungs of the colonized mice (**Figure 2B**). This is consistent with the amounts of LPS expected to transit from the gut microbiome to the lung as part of the gut-lung axis to establish along with associated SCFA levels, the resting lung immune tone. Finally, we used a *B.theta* mutant (del propionate) that was unable to produce C3/propionate to demonstrate that propionate from the intestine could travel to lung tissue: colonization by gavage of GF mice with C3 mutant *B.theta* resulted in less propionate present in the lungs of mice colonized compared to colonization with wildtype *B.theta* (**Figure 2C**).

**Figure 2.**
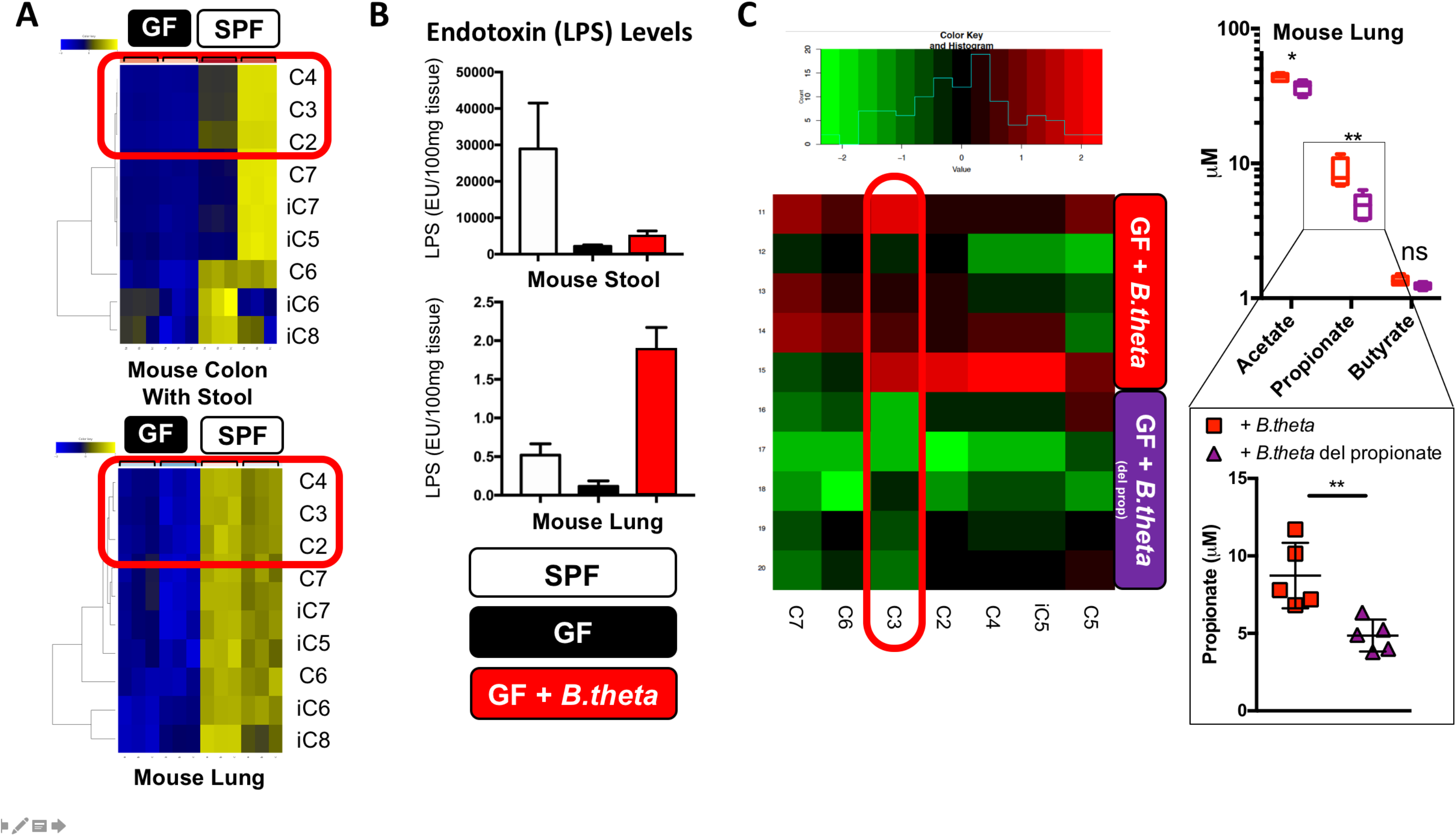
Mouse lung tissue SCFA and LPS levels depend on the presence of a gut microbiome that is metabolically active. (A) Profiling of SCFAs from SPF and germ free (GF) mice (n=2 littermates, and three independent colon/stool and lung tissue samples). C2-4 are highlighted by the red box. (B) Endotoxin (LPS) levels measured in mouse stool and lungs from SPF, GF and GF mice colonized with WT *B.theta* (~10^9^ CFU). (C) GF mice were colonized with *B.*theta: either WT (+*B.theta*) or a propionate-production deficient mutant (+*B.theta* del propionate) (~10^9^ CFU) and SCFA levels were profiled (left). C3 is highlighted by the red box. Measured levels of C2-4 are shown (right top) with a focus on propionate levels (right bottom). Experiments were repeated 2 times.

### SCFAs can alter mouse alveolar macrophage metabolism and responses to LPS exposure

We had previously reported that SCFA could significantly modulate AM response to LPS (or lipoteichoic acid) (50). Here, we wanted to understand the effects of SCFA and LPS co-presence on lung AM metabolism using Seahorse™ extracellular flux analyzer. We examined the metabolic response (glycolysis and oxidative phosphorylation) of AM to LPS and the effects of co-exposure to SCFA (**Figure 3A**, **top**). We observed that while low dose LPS exposure reduced maximal respiration and spare respiratory capacity of AM (blue bars), high dose propionate/C3 (purple bars) could partially reverse these LPS effects (**Figure 3A**, **bottom left and right**). Next, AMs were subjected to metabolic stress tests and monitored using Seahorse™. We found that AMs exposed to C3 displayed aerobic and energetic metabolic profiles, while LPS challenged AM were more glycolytic (**Figure 3B**). High dose C3, as we noted earlier, was able to partially reverse the metabolic phenotype of LPS-exposed AM (**Figure 3B**). In **Figure 3C** we present the metabolic profiles of AM co-exposed to no LPS and high dose C3 (↑C3) versus those exposed to LPS and low dose C3 (↓C3). As we have previously reported, the former conditions mimic the predicted antibiotic-treated lung environment which resulted in reduced lung inflammation following sterile injury and the significant enrichment of gram-positive propionate-producing *Lachnospiraceae* (51), while the latter conditions mimic the control group with an unperturbed gut microbiome and in whom robust lung inflammation ensues after sterile ischemia-reperfusion (IR) lung injury. Conditions that seemed to favor more inflammation correlated with AM metabolic profiles with lesser metabolic reserve, reduced ability to increase oxygen consumption rate when stressed, and greater extra cellular acidification rate (**Figure 3C**, right panels).

**Figure 3.**
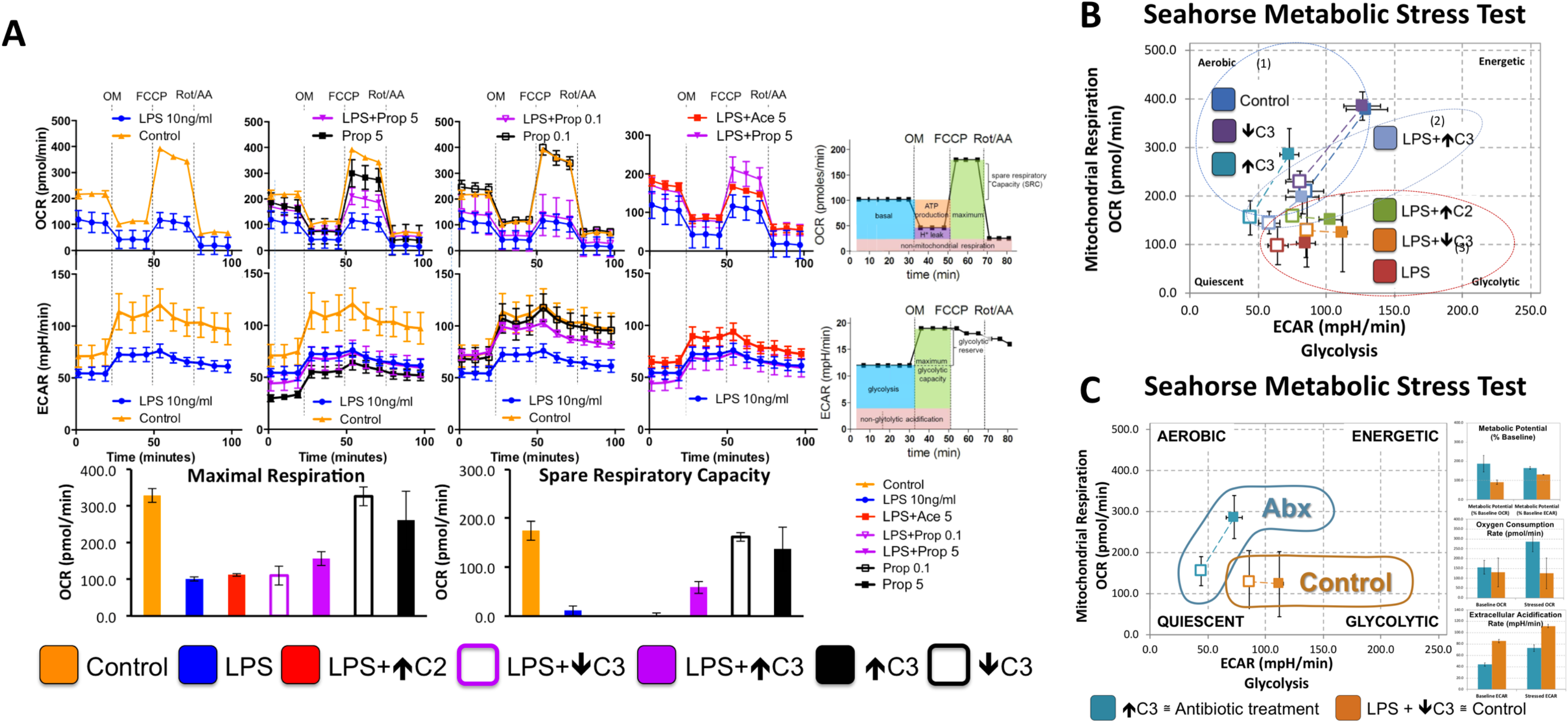
Alveolar macrophage immunometabolic response to LPS can be modulated by SCFAs. (A) AM cell line (MH-S) was unstimulated (control) or challenged with LPS (10 ng/ml), 0.1 mM propionate (↓C3), 5 mM propionate (↑C3), LPS plus 5 mM acetate (LPS +↑C2), LPS plus 0.1 mM propionate (LPS + ↓C3), LPS plus 5 mM propionate (LPS + ↑C3) after seeding cells 4 hours. Oxygen consumption rate (OCR) and extracellular acidification rage (ECAR) were measured under four blocks corresponding each effector injection, basal, ligomycin (OM), FCCP and rotenone+antimycin-a (Rot/AA) by using agilent Seahorse XFe24 analyzer platform. (B) Metabolic stress tests were run to measure mitochondrial respiration to compare metabolic phenotypes and profiles. Three circled groups represent three different metabolic stressed profiles: (1) control, ↓C3, and ↑C3; (2) LPS+↑C3; and (3) LPS, LPS+↓C3, and LPS+↑C2. (C) Two conditions were contrasted: exposure to a) ↑C3 alone and b) LPS+↓C3. All data was standardized to total protein using bicinchoninic acid assay (BCA). Experiments were repeated 2 times.

### Specific subsets of naïve and LPS stimulated mouse AM subsets express IL-1β

In order to further understand the effect of the gut microbiome on lung immune tone, we examined the baseline and LPS-stimulated expression of IL-1β within the mouse lung. Using spatial transcriptomic RNA probe pools (RNAScope™) to detect IL-1β and surfactant C mRNA, we had previously showed that resting, LPS-induced, and lung IR-induced IL-1β mRNA was expressed along the alveolar surface and in close proximity to AT2 cells and that IL-1β was not expressed by AT1 (HopX expressing) cells both in naïve and LPS-challenged lungs (51). Some CD11c+ AMs residing within the alveolar space appeared to express the naïve lung IL-1β signal indicating that a subset of AM may be responsible for IL-1β-dependent lung immune tone – both in the naïve and LPS-induced lungs (**Figure 4A**). These IL-1β expressing cells were confirmed to be CD45+ (hematopoietically-derived) (**Figure 4B**). Further imaging here revealed punctate IL-1β mRNA expression in the alveolar space appeared in close conjunction to AT2 cells (Surfactant C expressing) cells (**Figure 4c**) suggesting some type of coordination between AT2 and AMs with regard to IL-1β lung immune tone/priming. We also probed these lungs for IL-18, a member of the IL-1β family of cytokines also regulated by the inflammasome. IL-18 has been shown to be important for various lung disease pathologies (1, 26, 36, 58, 59), and could also serve as a potential contributor to lung immune tone. Interestingly, we found that IL-18 was expressed in a separate subset of cells distinct from IL-1β expressing cell (**Figure 4D**) suggesting perhaps of the existence of IL-1β and IL-18 specialized AM subsets.

**Figure 4.**
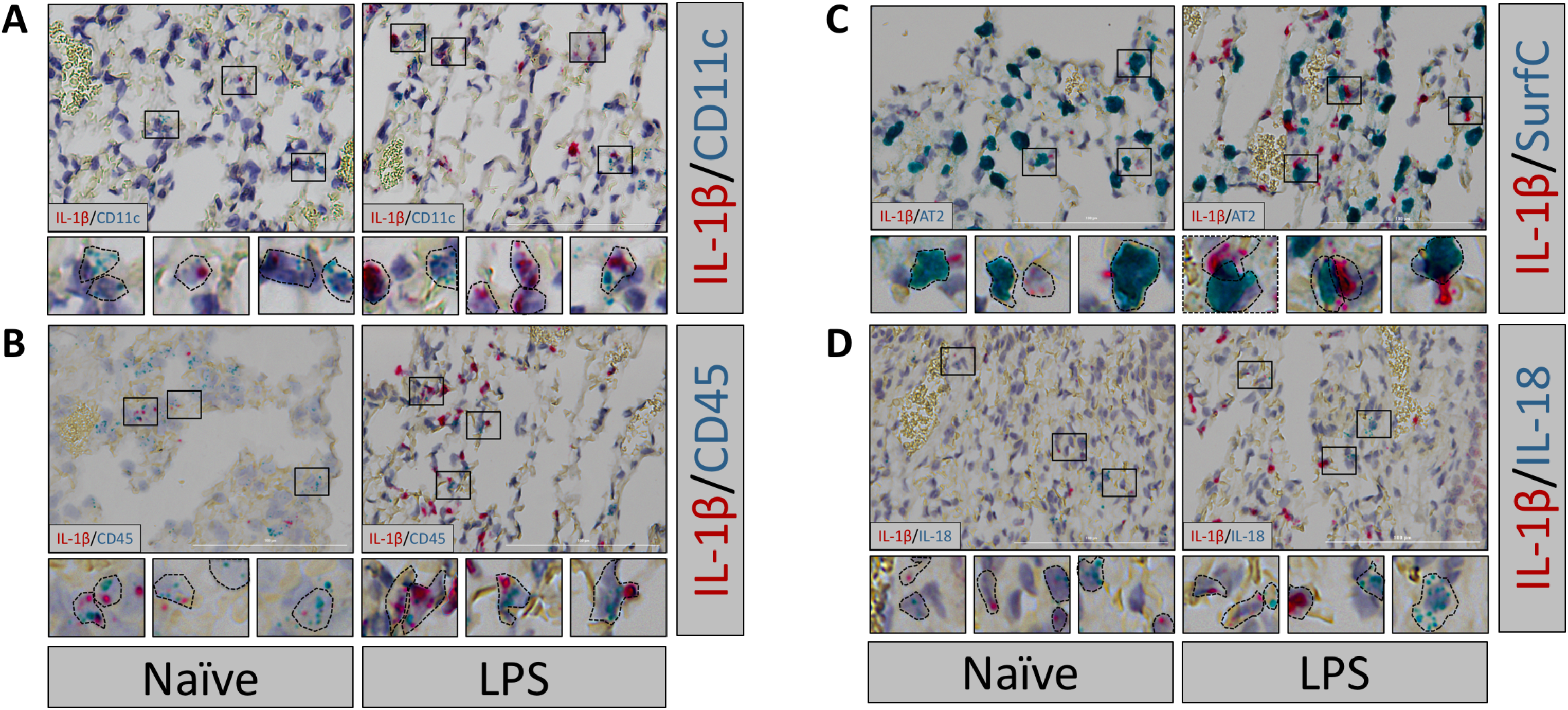
IL-1β presence in naïve mouse AM subsets and increased expression after LPS. Naïve wild-type C3H mice were treated with LPS (10 mg/kg IV for 1h). Lungs were harvested, fixed and stained using RNAscope® (from ACD) using specific mouse RNA probes for IL1β in combination with (A) CD11c (AM and dendritic cells), (B) CD45 (hematopoietic cells), (C) surfactant C (SurfC; AT 2 cells), and (D) IL-18. Dashed lines represent approximate estimations of cellular outlines of positively staining cells.

### FFAR2 and FFAR3 are expressed in AT2 and AM mouse cell lines and are inducible by LPS

How lungs sense signals transmitted from the gut is not well understood. We focused on mouse lung cell lines, such as AT2 (MLE-12 cells), AM (MH-S), and EC (EOMA) and first examined the LPS responsiveness of these cell lines *in vitro* and found that the EC more than AM were the major sources of IL-6 production (**Figure 5A**). Baseline expression of IL-1β and FFAR2 were highest in AM, and FFAR3 was similarly expressed in AT2 and AM (**Figure 5B**). Low dose LPS (10 ng/mL) resulted in a large increase in AM’s IL-1β, and AT2’s FFAR2 and FFAR3 expression (**Figure 5C**). Higher dose LPS (100 ng/mL) further increased the AM’s IL-1β and FFAR2 expression, and AT2’s FFAR3 expression (**Figure 5C**). Therefore, low level (such as may be seen in naïve mice) and high-level LPS (such as which could be seen in an infection or dysbiosis) resulted in very different profiles of FFAR2 and FFAR3 expression in alveolar resident cell types *in vitro*.

**Figure 5.**
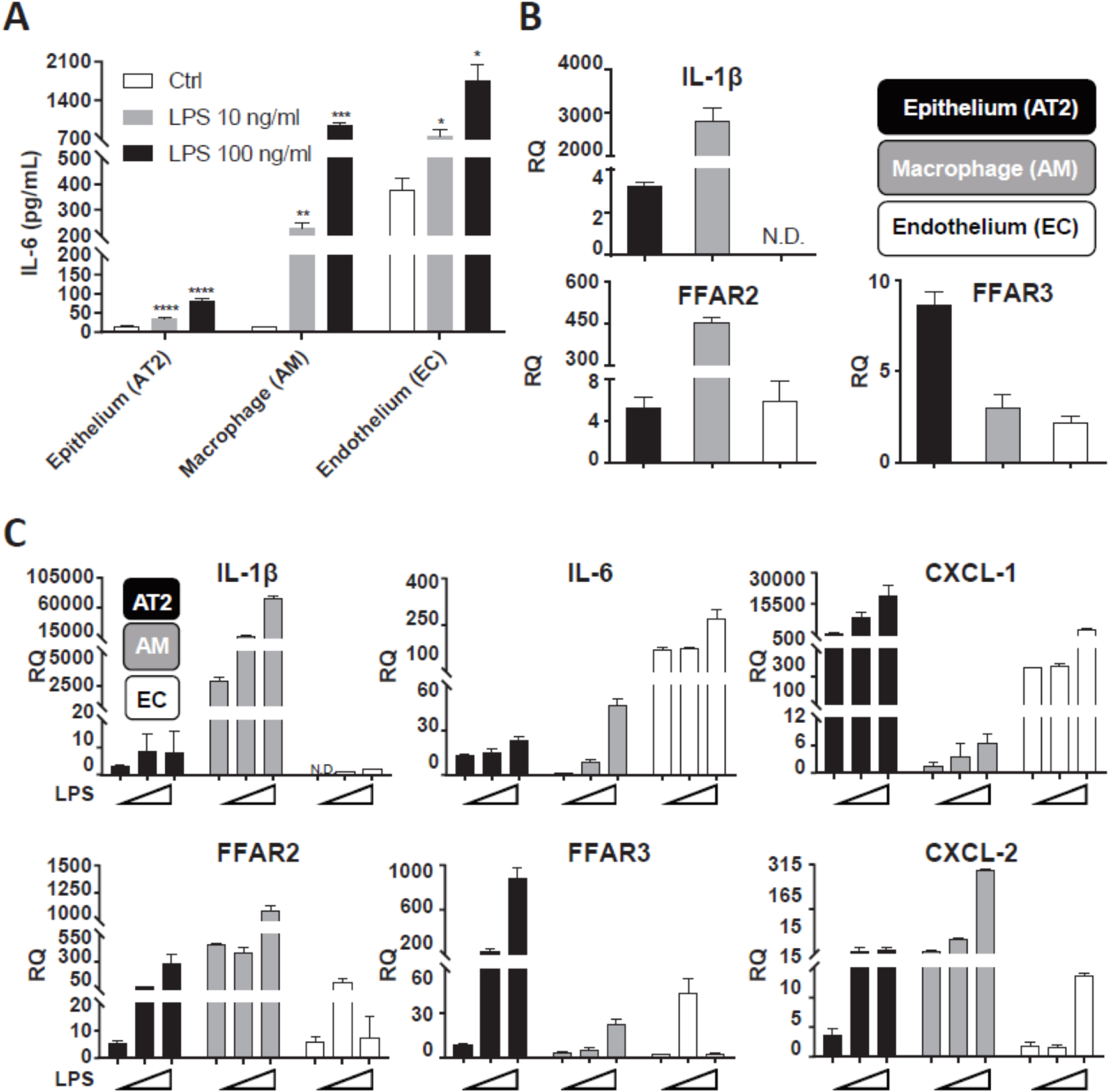
IL-1β is made by mouse AMs and FFAR2 and FFAR3 are expressed by AT2 and AMs but differentially. MLE 12 (mouse lung epithelial AT2), MH-S (wild-type BALB/c alveolar macrophage cell line AM), EOMA (129 background endothelial cell line EC) were treated with 10 ng/mL or 100 ng/mL LPS for 24 hours. Concentrations of IL-6 in cell culture supernatant were determined using the mouse DuoSet kit (A). Total RNA was extracted and expression of IL-1β, FFAR2, FFAR3, IL-6, CXCL-1, CXCL-2 was determined using RT-PCR (B, C). Experiments were repeated 3 times.

### Human alveolar cell expression of IL-1β and FFAR2 and FFAR3 ex vivo vs. in vitro culture

We measured *in vitro*/*ex vivo* expression of IL-1β and FFAR2 and FFAR3 in human BAL cells (AM) obtained from rejected human donor lungs and found that cells in culture overnight expressed more IL-1β and FFAR2 and less FFAR3 mRNA; and LPS exposure *in vitro* led to large and significant increases in all three gene’s expression levels (**Figure 6A**). Comparing freshly isolated AT2 cells, BAL AM, ECs from male and female cadaveric lungs, and HUVEC, we noted that AT2 and AMs were the primary source of IL-1β, FFAR2, and FFAR3 mRNA and not EC *in vitro* (**Figure 6B**). Interestingly, freshly isolated AT2 (versus the same cells after overnight culture in vitro) expressed more IL-1β message, less IL-6 and consistently low TLR4; in contrast cultured AM expressed more IL-1β, IL-6 and TLR4 versus freshly isolated BAL AM (**Figure 6C**).

**Figure 6.**
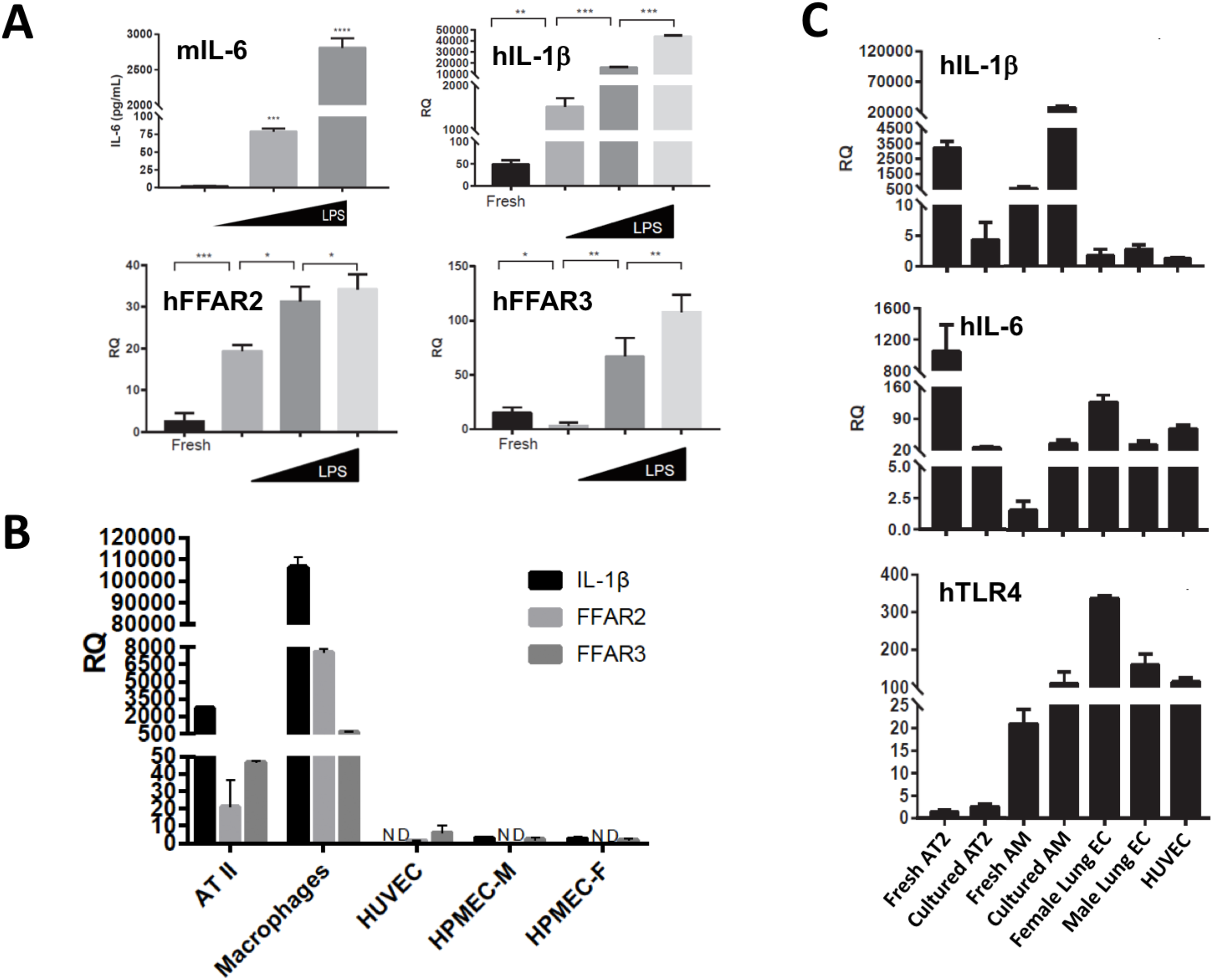
Human AMs express IL-1β and FFAR2 and FFAR3. Rejected human donor lung underwent bronchoalveolar lavage (BAL) to collect alveolar macrophages (AMs) or alveolar type II (AT2) cells were isolated from elastase-digested human lung tissue (from a different rejected human donor lung) using negative selection (anti-CD14 and IgG antibodies). (A) BAL AMs were assayed without further treatment (fresh) or treated with 0 ng/ml (control), 10 ng/mL or 100 ng/mL LPS for 24h. (B) Isolated AT2 cells (Fresh AT2), BAL AMs (Fresh AM), and commercially obtained primary human pulmonary microvascular endothelial cells - male or female (Female or Male Lung EC) or primary human umbilical vein endothelial cells (HUVEC) were assayed for expression of IL-1β, FFAR2 and FFAR3 by RT-qPCR. (C) Human BAL AMs and AT2 – either freshly isolated or after 24h in culture, female and male lung EC, and HUVEC were assayed for IL-1β (Hu IL-1β), IL-6 (Hu IL-6), and TLR-4 (Hu TLR4) expression by RT-qPCR.

### Human cellular expression of IL-1β and FFAR2 and FFAR3 in situ in fresh human lung tissue and BAL AMs

Our previous results suggested that alveolar cells outside of their natural lung microenvironment context may have very different expression of IL-1β, FFAR2, and FFAR3 and perhaps other genes. Therefore, we examined expression of these genes along with cell-specific markers *in situ* in the human lung. We noted that IL-1β signal in naïve human lungs was present in a subset of AMs that were CD11c/ITGAX positive (**Figure 7A**) and CD206/MRC1 positive (**Figure 7B**). Once again, we noted the close proximity of naïve lung IL-1β mRNA signal to AT2 cells (SFTPC/surfactant C positive (**Figure 7A**, far right inset b). FFAR2 and FFAR3 were co-expressed in MRC1/CD206 positive AMs as well as in other yet unidentified cell types (**Figure 7C**) but not by AT2 cells (**Figure 7D**). We exposed sections of the human lung tissue to LPS for 2h to stimulate the expression of IL-1β and found that a subset of AMs which were ITGAX/CD11c positive and CD206/MRC1 positive expressed IL-1β both at baseline and after LPS challenge (**Figure 7E-F**). To assess the level of variability in human lung immune tone, we assayed two different donor lung BAL cells for IL-1β, FFAR2 and FFAR3 expression and found 2-4 orders of magnitude differences in all three genes mRNA (**Figure 7G**). Moreover, the relative levels of these three genes seemed to change similarly implying a possible coordinated expression of this gene set.

**Figure 7.**
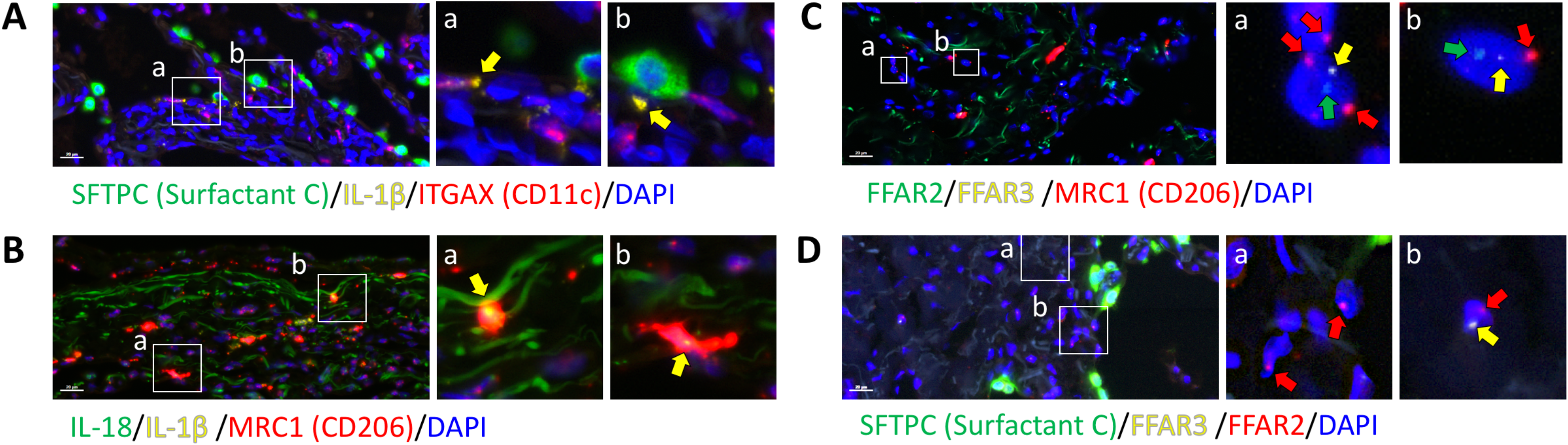
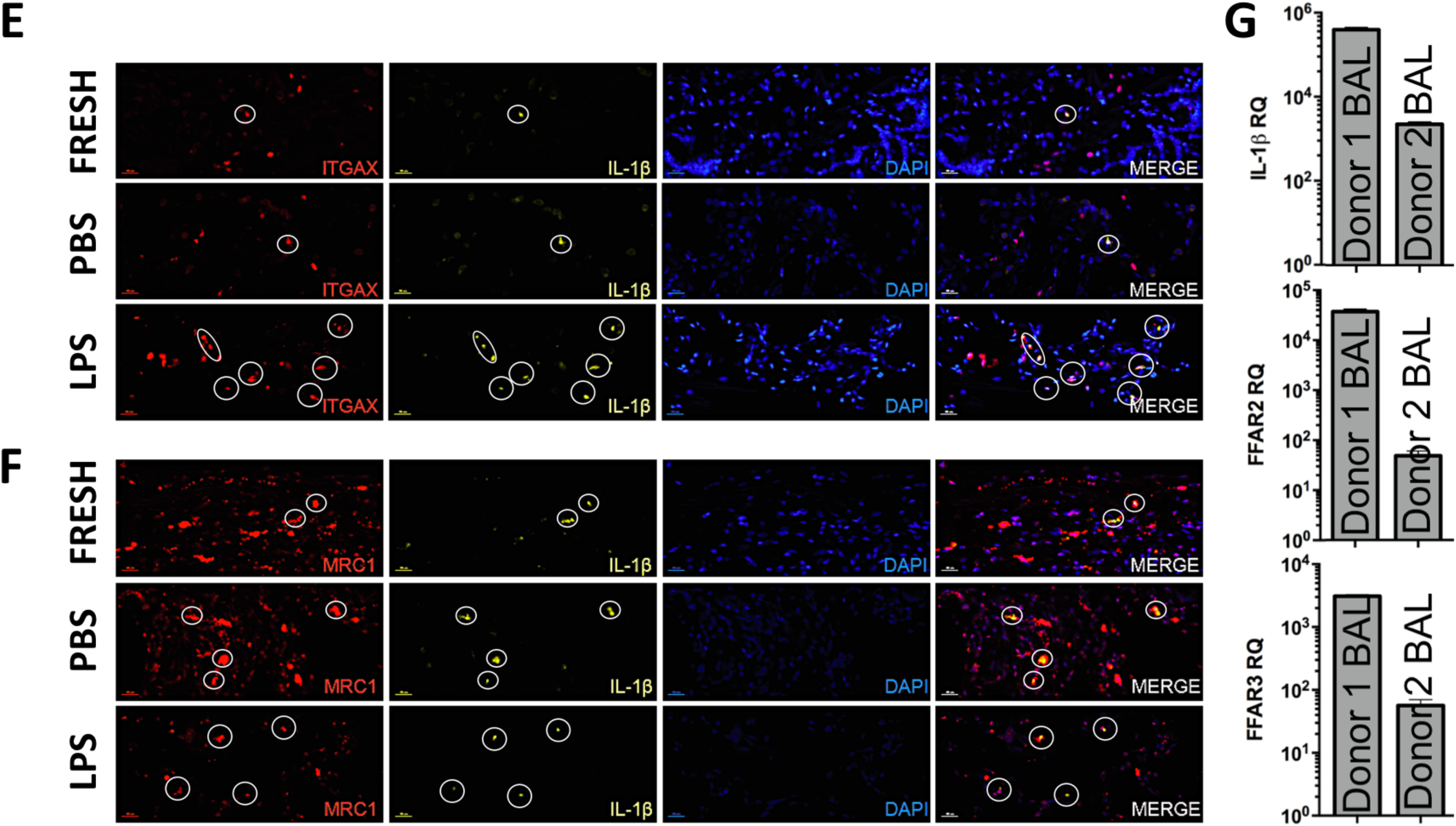
Human IL-1β, FFAR2 and FFAR3 are expressed in AMs *in vivo*. (A-C) Human lung sections were either fixed in formalin immediately after excision from patient and then stained/probed with DAPI and combinations of (A) surfactant C (AT2 cells, green), IL-1β (yellow), and ITGAX/CD11c (AM and dendritic cells, red); or (B) IL-18 (green), IL-1β (yellow), and CD206/MRC1 (macrophages, red); or (C) FFAR2 (green), FFAR3 (yellow), MRC1/CD206 (macrophages, red); or (D) SFTPC (Surfactant C, green), FFAR2 (yellow) and FFAR3 (red); (E & F) Human lungs were fixed immediately (FRESH) or incubated with PBS or LPS for 2h prior to fixation and then stained/probed with DAPI and (E) ITGAX/CD11c (red channel), and IL-1β (yellow channel); or (F) MRC1/CD206 (red channel) and IL-1β (yellow channel). (G) Human IL-1β, FFAR2, and FFAR3 mRNA expression were determined by RT-qPCR in freshly isolated BAL alveolar macrophages (AMs) from two different rejected human donor lungs.

## DISCUSSION

The major conclusions of this study are that human lungs contain metabolites and other bacterial signals that originate from the intestine, and that these signals likely contribute to the immune tone of the lung, namely the resting readiness state that the lungs exist in prior to injury or infection. These conclusions are based on the following experimental evidence. First, we demonstrated SCFA metabolites exist in human lung tissue and confirmed in mice that lung SCFA levels depended on intact and functional gut microbiota. Next, we demonstrated that the metabolic effects of LPS exposure on lung macrophages was profoundly affected by the co-presence of SCFAs. We then examined naïve and injured mouse lungs and found clear presence of immune tone via IL-1β expression within alveolar cells, as well as expression of SCFA receptors, namely FFAR2 and FFAR3, in alveolar macrophages (AM) and alveolar type 2 epithelial (AT2) cells. Exposure to LPS also regulated the expression of FFAR2 and FFAR3. Confirming these findings in human lung cells, we detected IL-1β, FFAR2 and FFAR3 expression in alveolar cells *in vitro* and *in vivo*. Collectively, this experimental evidence strongly supports the concept of lung immune tone as defined by the basal expression of pro-IL-1β message (Inflammasome signal 1), which is regulated by the gut microbiome through transport of bacterial products, including metabolites and LPS, along the gut-lung immune axis.

We previously demonstrated through the use of antibiotics in mice that the gut microbiome regulates sterile lung inflammation (42). Furthermore, we reported that IL-1β and its downstream regulation by the NLRP3 inflammasome was required for both sterile inflammatory and infectious immune responses (51). Finally, we uncovered a connection between propionate-producing gut bacteria and reduced sterile lung inflammation and observed that direct propionate introduction into the lung reduced lung anti-bacterial defense (50). Other groups have also described the gut-lung axis [reviewed in (7, 11)]. The Marsland and Lynch groups have published seminal work suggesting strong indirect and direct effects of the gut microbiome on allergic asthma in mice and humans (17, 31, 33, 53). Our study focused on the direct role of metabolites on influencing lung inflammation (18, 31). FFAR2 and FFAR3 have recently been reported in multiple cells in lung tissue including airway smooth muscle (35), airway epithelium (23, 54), mesenchymal cells (44), neutrophils, and alveolar macrophages (19). We speculate that immune, metabolic, and other regional heterogeneity within each lung may reflect the need for the lung to compartmentalize its various functions and to allow gas exchange, inflammation generation and resolution, and housekeeping/regeneration to occur simultaneously and this has been supported by recent reports (20, 56). Finally, that SCFAs can contribute to immune tone is supported in the literature by reports that SCFAs have pro-inflammatory effects (27), that LPS up-regulates FFAR2 and FFAR3 (2), and that acetate (via FFAR2) affects type I interferon responses after RSV infection (3).

We chose to focus on the gut microbiome and not the lung/airway microbiome based on the high metabolic activity that would be required for SCFA production. The distal ileum and colon contain a million to billion fold more bacteria per gram of organ weight in the distal ileum and colon as compared to the healthy lung (45). Many groups have strongly supported the role of the lung microbiome in influencing ARDS, sepsis and other lung disease outcomes (4, 12, 13, 38). This may be due to a greater microbial biomass within context of diseased lungs (where favorable growth conditions may exist) and due to gut bacterial translocation to the lung in critical illness [reviewed in (32, 55)]. Additionally, the effects of the gut microbiome and SCFAs on T_reg_ development and dendritic cell function locally in the colon, and remotely in the lung, has been shown to also contribute to the gut-lung axis (48, 53, 57, 60). Our experiments examined baseline steady-state immune tone in healthy lungs and immediate injury responses when gut-originating SCFAs likely have the strongest influence on early lung inflammatory responses. Also, while we focused in this study on SCFAs signaling via FFARs, attention should be also given to the suppressive effects of histone deacetylase (HDAC) inhibition on how propionate and butyrate mediate inflammatory responses (9). Notably, HDAC inhibition requires millimolar concentrations of SCFA usually seen in the colon (vs micromolar lung SCFA levels). Additionally, FFAR2-independent effects of SCFAs have been reported in lung eosinophils and mast cells in asthma and allergic airway disease (16, 49).

The concept of lung immune tone may be important to consider in field of lung biology. In the past, it was understood that as people age, their lungs were exposed to numerous potentially injurious factors and with each exposure/injury, the lung may heal and/or assume a “new normal” state/immune tone which would in turn influence the response to subsequent damage. The idea that the gut microbiome produces factors that signal to distant barrier organs such as the lung, to influence its health and host defense capability is very intriguing, as a means to understand varied responses to similar forms of lung injury. In severe systemic disease, the phenomenon of the intestinal barrier disruption or ‘leaky gut’ is well known but the influence of intestinal contents in health is less familiar. In fact, the immune tone of other organs, such as the placenta, has been proposed to be influenced by gut microbiome-derived SCFAs and LPS. Dissecting the rheostat-like dynamic control of lung inflammatory and immune responses could lead to better understanding of this baseline lung immune tone.

Our study has limitations: we focused on the two major SCFA receptors, namely FFAR2 and FFAR3, and not on others, such as GPR109a and OLFR78 (10). GPR109a primarily resides in the gut where its primary ligand (butyrate) is present (25), and the olfactory chemo sensor OLFR78, which binds propionate, acetate and lactate, is found in the upper airway only (8, 10, 39, 52). Other metabolites, such as 12,13-dihydroxy-9Z-octadecenoic acid (12, 13-diHOME) and others including primary and secondary bile acids, may also participate in the gut-lung immune axis and establishment of lung immune tone (31). Another limitation is the lack of definitive findings that interrupting the gut-lung immune axis and lung sensing of SCFAs has clear effect on lung inflammatory and immune responses. Some of these findings have been reported by others, specifically using global FFAR knockout mice and cells and studying bacterial pneumonia (19), RSV infection (3), or influenza infection (54). Further studies will require the generation of lung-specific FFAR knockout mice and careful inhibitor studies that are topics of future investigations in our laboratory. We have previously reported that exogenously short-circuiting this gut-lung immune axis by providing propionate directly to mouse lungs can strongly influence lung immunity to bacterial pneumonia (50) and further experiments to understand the dynamic and mutable nature of lung immune tone will be needed.

The major conclusions of this study are that the lung possesses the capability to sense SCFA metabolites by extension the gut microbiome, can influence and regulate lung immune tone, namely how the lung responds to injury and infection. Moreover, this effect on immune tone may be driven by changes in cellular metabolism as a result of local SCFA concentrations. The implication of these findings are that diet, medication, and lifestyle influences on the gut microbiome may indirectly have substantial effects on lung inflammatory and immune responses. Moreover, these effects may be more amenable to beneficial intervention via specific SCFA supplementation, diets including fermentable fiber or fecal microbiota transplantation (FMT). Overall, the gut-lung immune axis, not unlike many other axes connecting the gut microbiome to vital organ systems, may be an avenue towards better understanding lung biology in health and disease as well as integrating known effects of genetics, diet and environment on gut and lung function.

## ACKNOWLEDGMENTS

We would like to acknowledge the following individuals for assistance with providing lung tissue, reagents, mice, advice, helpful discussions, and critical reading and editing of the manuscript: Arthur Hill (UCSF), Judith Hellman (UCSF), Michael Matthay (UCSF), Mervyn Maze (UCSF), Kevin Wilhelmsen (BioAge Labs), Eric Martens (University of Michigan). The authors would also like to acknowledge the continuing support of THFC (COYS) on these studies.

## REFERENCES

1. Abdel Fattah E, Bhattacharya A, Herron A, Safdar Z, Eissa NT. Critical role for IL-18 in spontaneous lung inflammation caused by autophagy deficiency. J Immunol 194: 5407–16, 2015.

2. Ang Z, Er JZ, Tan NS, Lu J, Liou YC, Grosse J, Ding JL. Human and mouse monocytes display distinct signalling and cytokine profiles upon stimulation with FFAR2/FFAR3 short-chain fatty acid receptor agonists. Sci Rep 6, 2016.

3. Antunes KH, Fachi JL, de Paula R, da Silva EF, Pral LP, dos Santos AÁ, Dias GBM, Vargas JE, Puga R, Mayer FQ, Maito F, Zárate-Bladés CR, Ajami NJ, Sant’Ana MR, Candreva T, Rodrigues HG, Schmiele M, Silva Clerici MTP, Proença-Modena JL, Vieira AT, Mackay CR, Mansur D, Caballero MT, Marzec J, Li J, Wang X, Bell D, Polack FP, Kleeberger SR, Stein RT, Vinolo MAR, de Souza APD. Microbiota-derived acetate protects against respiratory syncytial virus infection through a GPR43-type 1 interferon response. Nat Commun 10, 2019.

4. Barcik W, Boutin RCT, Sokolowska M, Finlay BB. The Role of Lung and Gut Microbiota in the Pathology of Asthma. Immunity 52Cell Press.: 241–255, 2020.

5. Bonini JA, Anderson SM, Steiner DF. Molecular cloning and tissue expression of a novel orphan G protein-coupled receptor from rat lung. Biochem Biophys Res Commun 234: 190–193, 1997.

6. Brown AJ, Goldsworthy SM, Barnes AA, Eilert MM, Tcheang L, Daniels D, Muir AI, Wigglesworth MJ, Kinghorn I, Fraser NJ, Pike NB, Strum JC, Steplewski KM, Murdock PR, Holder JC, Marshall FH, Szekeres PG, Wilson S, Ignar DM, Foord SM, Wise A, Dowell SJ. The orphan G protein-coupled receptors GPR41 and GPR43 are activated by propionate and other short chain carboxylic acids. J Biol Chem 278: 11312–11319, 2003.

7. Budden KF, Gellatly SL, Wood DLA, Cooper MA, Morrison M, Hugenholtz P, Hansbro PM. Emerging pathogenic links between microbiota and the gut–lung axis. Nat Rev Microbiol 15: 55–63, 2017.

8. Chang AJ, Ortega FE, Riegler J, Madison D V., Krasnow MA. Oxygen regulation of breathing through an olfactory receptor activated by lactate. Nature 527: 240–244, 2015.

9. Chang P V., Hao L, Offermanns S, Medzhitov R. The microbial metabolite butyrate regulates intestinal macrophage function via histone deacetylase inhibition. Proc Natl Acad Sci 111: 2247–2252, 2014.

10. Corrêa-Oliveira R, Fachi JL, Vieira A, Sato FT, Vinolo MAR. Regulation of immune cell function by short-chain fatty acids. Clin Transl Immunol 5: e73, 2016.

11. Dang AT, Marsland BJ. Microbes, metabolites, and the gut–lung axis. Mucosal Immunol. 12Nature Publishing Group.: 843–850, 2019.

12. Dickson RP. The lung microbiome and ARDS it is time to broaden the model. Am. J. Respir. Crit. Care Med. 197American Thoracic Society.: 549–551, 2018.

13. Dickson RP, Erb-Downward JR, Martinez FJ, Huffnagle GB. The Microbiome and the Respiratory Tract. Annu Rev Physiol 78: 481–504, 2016.

14. Dickson RP, Huffnagle GB. The Lung Microbiome: New Principles for Respiratory Bacteriology in Health and Disease. PLoS Pathog 11: e1004923, 2015.

15. Fang X, Neyrinck AP, Matthay MA, Lee JW. Allogeneic human mesenchymal stem cells restore epithelial protein permeability in cultured human alveolar type II cells by secretion of angiopoietin-1. J Biol Chem 285: 26211–26222, 2010.

16. Folkerts J, Redegeld F, Folkerts G, Blokhuis B, van den Berg MPM, de Bruijn MJW, van IJcken WFJ, Junt T, Tam SY, Galli SJ, Hendriks RW, Stadhouders R, Maurer M. Butyrate inhibits human mast cell activation via epigenetic regulation of Fc∊RI-mediated signaling. Allergy Eur. J. Allergy Clin. Immunol. (2020). doi: 10.1111/all.14254.

17. Fujimura KE, Demoor T, Rauch M, Faruqi AA, Jang S, Johnson CC, Boushey HA, Zoratti E, Ownby D, Lukacs NW, Lynch S V. House dust exposure mediates gut microbiome Lactobacillus enrichment and airway immune defense against allergens and virus infection. Proc Natl Acad Sci U S A 111: 805–810, 2014.

18. Fujimura KE, Sitarik AR, Havstad S, Lin DL, Levan S, Fadrosh D, Panzer AR, LaMere B, Rackaityte E, Lukacs NW, Wegienka G, Boushey HA, Ownby DR, Zoratti EM, Levin AM, Johnson CC, Lynch S V. Neonatal gut microbiota associates with childhood multisensitized atopy and T cell differentiation. Nat Med 22: 1187–1191, 2016.

19. Galvão I, Tavares LP, Corrêa RO, Fachi JL, Rocha VM, Rungue M, Garcia CC, Cassali G, Ferreira CM, Martins FS, Oliveira SC, Mackay CR, Teixeira MM, Vinolo MAR, Vieira AT. The metabolic sensor GPR43 receptor plays a role in the control of Klebsiella pneumoniae infection in the lung. Front Immunol 9, 2018.

20. Garg N, Wang M, Hyde E, da Silva RR, Melnik A V., Protsyuk I, Bouslimani A, Lim YW, Wong R, Humphrey G, Ackermann G, Spivey T, Brouha SS, Bandeira N, Lin GY, Rohwer F, Conrad DJ, Alexandrov T, Knight R, Dorrestein PC. Three-Dimensional Microbiome and Metabolome Cartography of a Diseased Human Lung. Cell Host Microbe 22: 705–716.e4, 2017.

21. Ghorbani P, Santhakumar P, Hu Q, Djiadeu P, Wolever TMS, Palaniyar N, Grasemann H. Short-chain fatty acids affect cystic fibrosis airway inflammation and bacterial growth. Eur Respir J 46: 1033–1045, 2015.

22. Gollwitzer ES, Marsland BJ. Microbiota abnormalities in inflammatory airway diseases - Potential for therapy. Pharmacol. Ther. 141Pharmacol Ther.: 32–39, 2014.

23. Imoto Y, Kato A, Takabayashi T, Sakashita M, Norton JE, Suh LA, Carter RG, Weibman AR, Hulse KE, Stevens W, Harris KE, Peters AT, Grammer LC, Tan BK, Welch K, Conley DB, Kern RC, Fujieda S, Schleimer RP. Short-chain fatty acids induce tissue plasminogen activator in airway epithelial cells via GPR41&43. Clin Exp Allergy 48: 544–554, 2018.

24. Jacobson A, Lam L, Rajendram M, Tamburini F, Honeycutt J, Pham T, Van Treuren W, Pruss K, Stabler SR, Lugo K, Bouley DM, Vilches-Moure JG, Smith M, Sonnenburg JL, Bhatt AS, Huang KC, Monack D. A Gut Commensal-Produced Metabolite Mediates Colonization Resistance to Salmonella Infection. Cell Host Microbe 24: 296–307.e7, 2018.

25. Jobin C. GPR109a: The missing link between microbiome and good health? Immunity 40NIH Public Access.: 8–10, 2014.

26. Jordan JA, Guo RF, Yun EC, Sarma V, Warner RL, Crouch LD, Senaldi G, Ulich TR, Ward PA. Role of IL-18 in acute lung inflammation. J Immunol 167: 7060–8, 2001.

27. Kim MH, Kang SG, Park JH, Yanagisawa M, Kim CH. Short-chain fatty acids activate GPR41 and GPR43 on intestinal epithelial cells to promote inflammatory responses in mice. Gastroenterology 145, 2013.

28. Kimura I, Ichimura A, Ohue-Kitano R, Igarashi M. Free fatty acid receptors in health and disease. Physiol Rev 100: 171–210, 2020.

29. Korpela K, Salonen A, Virta LJ, Kekkonen RA, Forslund K, Bork P, De Vos WM. Intestinal microbiome is related to lifetime antibiotic use in Finnish pre-school children. Nat Commun 7: 1–8, 2016.

30. Kovatcheva-Datchary P, Nilsson A, Akrami R, Lee YS, De Vadder F, Arora T, Hallen A, Martens E, Björck I, Bäckhed F. Dietary Fiber-Induced Improvement in Glucose Metabolism Is Associated with Increased Abundance of Prevotella. Cell Metab 22: 971–982, 2015.

31. Levan SR, Stamnes KA, Lin DL, Fujimura KE, Ownby DR, Zoratti EM, Boushey HA, Johnson CC, Lynch S V. Neonatal gut-microbiome-derived 12,13 DiHOME impedes tolerance and promotes childhood atopy and asthma. bioRxiv (April 30, 2018). doi: 10.1101/311704.

32. Man WH, De Steenhuijsen Piters WAA, Bogaert D. The microbiota of the respiratory tract: Gatekeeper to respiratory health. Nat. Rev. Microbiol. 15Nature Publishing Group.: 259–270, 2017.

33. Marsland BJ, Trompette A, Gollwitzer ES. The gut-lung axis in respiratory disease. In: Annals of the American Thoracic Society. 2015.

34. Melhem H, Kaya B, Ayata CK, Hruz P, Niess JH. Metabolite-Sensing G Protein-Coupled Receptors Connect the Diet-Microbiota-Metabolites Axis to Inflammatory Bowel Disease. Cells 8: 450, 2019.

35. Mizuta K, Sasaki H, Zhang Y, Matoba A, Emala CW. The short-chain free fatty acid receptor FFAR3 is expressed and potentiates contraction in human airway smooth muscle. Am J Physiol Lung Cell Mol Physiol 318: L1248–L1260, 2020.

36. Nakajima T, Owen CA. Interleukin-18: The Master Regulator Driving Destructive and Remodeling Processes in the Lungs of Patients with Chronic Obstructive Pulmonary Disease? Am J Respir Crit Care Med 185: 1137–1139, 2012.

37. Nilsson NE, Kotarsky K, Owman C, Olde B. Identification of a free fatty acid receptor, FFA2R, expressed on leukocytes and activated by short-chain fatty acids. Biochem Biophys Res Commun 303: 1047–1052, 2003.

38. O’Dwyer DN, Dickson RP, Moore BB. The Lung Microbiome, Immunity, and the Pathogenesis of Chronic Lung Disease. J Immunol 196: 4839–47, 2016.

39. Pluznick JL, Protzko RJ, Gevorgyan H, Peterlin Z, Sipos A, Han J, Brunet I, Wan LX, Rey F, Wang T, Firestein SJ, Yanagisawa M, Gordon JI, Eichmann A, Peti-Peterdi J, Caplan MJ. Olfactory receptor responding to gut microbiotaderived signals plays a role in renin secretion and blood pressure regulation. Proc Natl Acad Sci U S A 110: 4410–4415, 2013.

40. Le Poul E, Loison C, Struyf S, Springael JY, Lannoy V, Decobecq ME, Brezillon S, Dupriez V, Vassart G, Van Damme J, Parmentier M, Detheux M. Functional characterization of human receptors for short chain fatty acids and their role in polymorphonuclear cell activation. J Biol Chem 278: 25481–25489, 2003.

41. Prakash A, Sundar S V., Zhu YG, Tran A, Lee JW, Lowell C, Hellman J. Lung Ischemia-Reperfusion is a Sterile Inflammatory Process Influenced by Commensal Microbiota in Mice. Shock 44: 272–279, 2015.

42. Prakash A, Sundar S, Zhu Y-G, Tran A, Lee J-W, Lowell C, Hellman J. Lung Ischemia Reperfusion (IR) is a Sterile Inflammatory Process Influenced by Commensal Microbiota in Mice. Shock 44: 272–9, 2015.

43. Rooks MG, Garrett WS. Gut microbiota, metabolites and host immunity. 16: 341–352, 2016.

44. Rutting S, Xenaki D, Malouf M, Horvat JC, Wood LG, Hansbro PM, Oliver BG. Short-chain fatty acids increase tnfα-induced inflammation in primary human lung mesenchymal cells through the activation of p38 mapk. Am J Physiol - Lung Cell Mol Physiol 316: L157–L174, 2019.

45. Sender R, Fuchs S, Milo R. Revised Estimates for the Number of Human and Bacteria Cells in the Body. PLOS Biol 14: e1002533, 2016.

46. Shapiro H, Thaiss CA, Levy M, Elinav E. The cross talk between microbiota and the immune system: metabolites take center stage. Curr Opin Immunol 30: 54–62, 2014.

47. Singh N, Gurav A, Sivaprakasam S, Brady E, Padia R, Shi H, Thangaraju M, Prasad PD, Manicassamy S, Munn DH, Lee JR, Offermanns S, Ganapathy V. Activation of Gpr109a, receptor for niacin and the commensal metabolite butyrate, suppresses colonic inflammation and carcinogenesis. Immunity 40: 128–139, 2014.

48. Smith PM, Howitt MR, Panikov N, Michaud M, Gallini CA, Bohlooly-Y M, Glickman JN, Garrett WS. The microbial metabolites, short-chain fatty acids, regulate colonic T reg cell homeostasis. Science (80-) 341: 569–573, 2013.

49. Theiler A, Bärnthaler T, Platzer W, Richtig G, Peinhaupt M, Rittchen S, Kargl J, Ulven T, Marsh LM, Marsche G, Schuligoi R, Sturm EM, Heinemann A. Butyrate ameliorates allergic airway inflammation by limiting eosinophil trafficking and survival. J Allergy Clin Immunol 144: 764–776, 2019.

50. Tian X, Hellman J, Prakash A. Elevated Gut Microbiome-Derived Propionate Levels Are Associated With Reduced Sterile Lung Inflammation and Bacterial Immunity in Mice. Front Microbiol 10: 159, 2019.

51. Tian X, Sun H, Casbon A-J, Lim E, Francis KP, Hellman J, Prakash A. NLRP3 Inflammasome Mediates Dormant Neutrophil Recruitment following Sterile Lung Injury and Protects against Subsequent Bacterial Pneumonia in Mice. Front Immunol 8: 1337, 2017.

52. Torres-Torrelo H, Ortega-Sáenz P, Macías D, Omura M, Zhou T, Matsunami H, Johnson RS, Mombaerts P, López-Barneo J. The role of Olfr78 in the breathing circuit of mice. Nature 561: E33–E40, 2018.

53. Trompette A, Gollwitzer ES, Yadava K, Sichelstiel AK, Sprenger N, Ngom-Bru C, Blanchard C, Junt T, Nicod LP, Harris NL, Marsland BJ. Gut microbiota metabolism of dietary fiber influences allergic airway disease and hematopoiesis. Nat Med 20: 159–166, 2014.

54. Wang G, Jiang L, Wang J, Zhang J, Kong F, Li Q, Yan Y, Huang S, Zhao Y, Liang L, Li J, Sun N, Hu Y, Shi W, Deng G, Chen P, Liu L, Zeng X, Tian G, Bu Z, Chen H, Li C. The G Protein-Coupled Receptor FFAR2 Promotes Internalization during Influenza A Virus Entry. J Virol 94, 2019.

55. Wypych TP, Wickramasinghe LC, Marsland BJ. The influence of the microbiome on respiratory health. Nat. Immunol. 20Nature Publishing Group.: 1279–1290, 2019.

56. Xu-Vanpala S, Deerhake ME, Wheaton JD, Parker ME, Juvvadi PR, MacIver N, Ciofani M, Shinohara ML. Functional heterogeneity of alveolar macrophage population based on expression of CXCL2. Sci Immunol 5, 2020.

57. Yang W, Xiao Y, Huang X, Chen F, Sun M, Bilotta AJ, Xu L, Lu Y, Yao S, Zhao Q, Liu Z, Cong Y. Microbiota Metabolite Short-Chain Fatty Acids Facilitate Mucosal Adjuvant Activity of Cholera Toxin through GPR43. J Immunol 203: 282–292, 2019.

58. Yasuda K, Nakanishi K, Tsutsui H. Interleukin-18 in Health and Disease. Int J Mol Sci 20, 2019.

59. Yasuda K, Nakanishi K, Tsutsui H. Interleukin-18 in Health and Disease. Int J Mol Sci 20, 2019.

60. Zaiss MM, Rapin A, Lebon L, Dubey LK, Mosconi I, Sarter K, Piersigilli A, Menin L, Walker AW, Rougemont J, Paerewijck O, Geldhof P, McCoy KD, Macpherson AJ, Croese J, Giacomin PR, Loukas A, Junt T, Marsland BJ, Harris NL. The Intestinal Microbiota Contributes to the Ability of Helminths to Modulate Allergic Inflammation. Immunity 43: 998–1010, 2015.

